# Activity-dependent long-term potentiation of electrical synapses in the mammalian thalamus

**DOI:** 10.1101/570101

**Authors:** Brandon Fricker, Emily Heckman, Patrick C. Cunningham, Julie S. Haas

## Abstract

Activity-dependent changes of synapse strength have been extensively characterized at chemical synapses, but the relationship between physiological forms of activity and strength at electrical synapses remains poorly understood. For mammalian electrical synapses composed of hexomers of connexin36, physiological forms of neuronal activity in coupled pairs has thus far have only been linked to long-term depression; activity that results in strengthening of electrical synapses has not yet been identified. The thalamic reticular nucleus (TRN), a central brain area primarily connected by gap junctional (electrical) synapses, regulates cortical attention to the sensory surround. Bidirectional plasticity of electrical synapses may be a key mechanism underlying these processes in both healthy and diseased states. Here we show in electrically coupled TRN pairs that tonic spiking in one neuron results in long-term potentiation of electrical synapses between coupled pairs of TRN neurons. Potentiation is expressed asymmetrically, indicating that regulation of connectivity depends on the direction of use. Further, potentiation depends on calcium flux, and we thus propose a calcium-based activity rule for bidirectional plasticity of electrical synapse strength. Because electrical synapses dominate intra-TRN connectivity, these synapses and their modifications are key regulators of thalamic attention circuitry. More broadly, bidirectional modifications of electrical synapses are likely to be a widespread and powerful principle for ongoing, dynamic reorganization of neuronal circuitry across the brain.

**Summary:** Long-term potentiation results from spiking in one cell of an electrically coupled pair. Asymmetry of synapses increases following unidirectional activity. We suggest a calcium-based rule for electrical synapse plasticity.

## Introduction

The thalamic reticular nucleus (TRN) is a central brain region exclusively comprising inhibitory neurons that gate the bidirectional flow of information between thalamus and cortex, and ultimately regulate the cognitive process of attention (Halassa et al., 2014; Kimura, 2014; McAlonan et al., 2006; Sherman, 2016; Zikopoulos & Barbas, 2012). Like all thalamic neurons, TRN neurons fire action potentials in two modes (Contreras et al., 1992). Bursts, which feature a slow calcium spike from a T-type calcium current (Huguenard & Prince, 1992) crowned by a fast barrage of sodium spikes, dominate TRN activity during slow sleep rhythms (Crunelli et al., 2006; Destexhe et al., 1996; Huguenard & Prince, 1992; McCormick & Bal, 1997; Steriade et al., 1993), are a feature of *absence* epilepsy (Fuentealba & Steriade, 2005), and are reduced in schizophrenics (Ferrarelli & Tononi, 2011). Regular tonic spikes, in contrast, are prevalent during attentive behaviors (Pinault, 2004). Because the main mode of intra-TRN communication is its dense electrical synapses (Hou et al., 2016; Landisman et al., 2002), understanding how and when electrical synapses change in strength during these two modes of activity is key for understanding attention processes and rhythm generation.

In the mature mammalian brain, electrical synapses are composed of paired hexomers of connexin36 that pass ions and small molecules, and mainly couple inhibitory GABAergic neurons (Bennett & Zukin, 2004; Connors & Long, 2004; Galarreta & Hestrin, 2001). Electrical synapses contribute to synchrony in coupled networks (Chow & Kopell, 2000; Destexhe, 1998; Draguhn et al., 1998; Gutierrez et al., 2013; Haas & Landisman, 2012; Pernelle et al., 2018; Pfeuty et al., 2005; Wang & Rinzel, 1993; Whittington & Traub, 2003) and regulate timing of spikes in the neurons they couple (Haas, 2015; Pham & Haas, 2018, 2019). Despite their prevalence and function between spiking neurons across the brain, the effects of neuronal activity on connection strength remain sparsely characterized. A line of work in non-mammalian nervous systems has demonstrated the possibility for central electrical synapses to undergo plasticity (McMahon et al., 1989; Pereda & Faber, 1996; Welzel & Schuster, 2018; Yang et al., 1990). In the mammalian brain, synaptic input has been shown to modulate mammalian electrical synapses in the inferior olive (Lefler et al., 2014) in an NMDA-dependent manner (Mathy et al., 2014; Turecek et al., 2014), and tetanic glutamatergic input to coupled TRN neurons results in long-term depression (LTD)(Landisman & Connors, 2005). The impact of spiking in coupled neurons on electrical synapses is much less well understood. We previously showed that LTD of electrical synapses in TRN also follows bursting activity of coupled neurons (Haas et al., 2011) in a calcium-dependent manner (Sevetson et al., 2017), but a link between activity in coupled neurons and strengthening of the synapse has remained elusive.

Here we demonstrate, using dual whole-cell patch recordings, that long-term potentiation (LTP) of electrical synapses in the TRN follows low-frequency spiking in one of the cells. LTP is specific to single-cell activity, and depends on calcium influx. We show that the increase in coupling strength is expressed in an asymmetrical manner that depends on the direction of synapse use. Combined with results of calcium imaging during activity, our work leads us to propose a calcium-based activity dependent plasticity rule for electrical synapses.

## Results

Thalamic neurons fire action potentials (APs) in two modes: from hyperpolarized resting potentials, bursts of calcium spikes crowned by a quick sequence of sodium spikes; and from depolarized rests, regular (tonic) sodium spikes (Fig. 1A). Having previously established that bursting in coupled TRN neurons leads to long-term depression of the electrical synapse between them (Haas et al., 2011) as a result of large-conductance, T-channel mediated calcium influx during bursts (Sevetson et al., 2017), we reasoned that tonic spikes and smaller calcium influx might lead to potentiation of electrical synapse strength. In order to minimize activation of the low voltage-activated T-type calcium current that underlies bursts, we added the specific antagonist TTA-A2 (Kraus et al., 2010) (1 μM) to the ACSF bath solution. We applied steady current to raise the membrane potential of one cell of a coupled pair at or above −55 mV, again to minimize the T current, while maintaining its coupled neighbor at −70 mV. We measured electrical coupling in each direction separately (Fig. 1B). We then applied small-amplitude, long pulses of current to the depolarized cell to induce non-continuous tonic spiking for 5 minutes, resulting in a spiking frequency of 5 - 10 Hz (Fig. 1C). After 5 minutes of tonic spiking activity in one neuron, coupling conductance measured from the quiet cell into the active cell increased on average by 15.5% ± 2.3% (p_t_ = 0.004) and coupling coefficient in the same direction increased by 13.6 ± 2.2% (Fig. 1E-F; p_t_ = 0.01, n = 12 pairs). Coupling measured by current injection into the active cell, in contrast, did not change in conductance (−1.5 ± 0.5%, p_t_ = 0.78) while coefficients in this direction decreased by 7.6 ± 0.5% (p_t_ = 0.018). Input resistance in the quiet cell decreased by 5.5 ± 0.4% (Fig. 1G; p_t_ = 0.0005), while input resistance in the active cell was unchanged (p_t_ = 0.73; sign tests performed on changes in cc and Gc gave similar results). Input resistance and coupling coefficients are directly related; thus it is difficult to determine whether the decrease in R_in_ caused the apparent decrease in cc or *vice versa*, but this dependence is minimized by the calculation of G_C_, which demonstrated clear changes in only one direction in this experiment. LTP was sustained for more than 25 minutes after induction.

**Figure 1.**
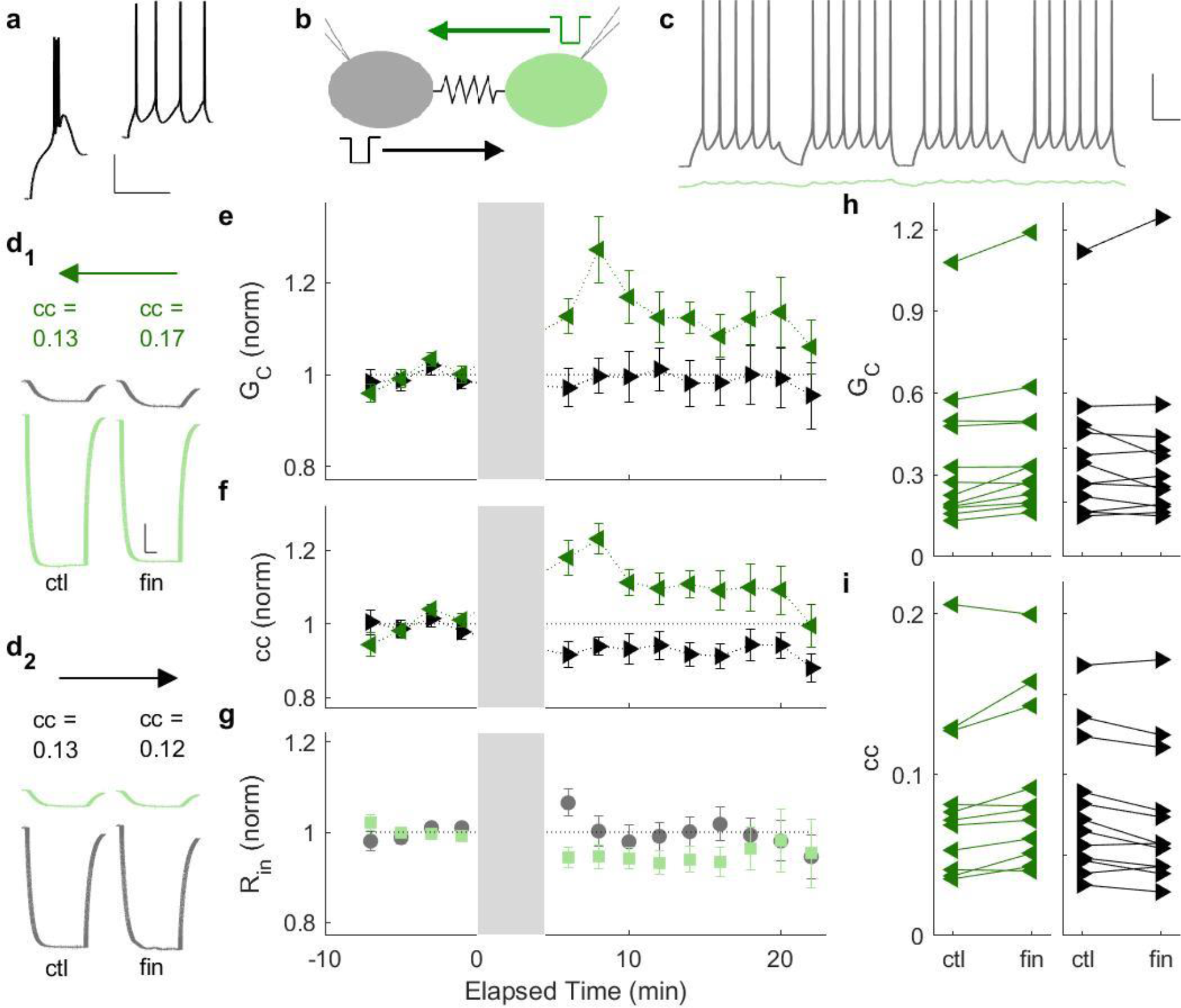
Long-term potentiation results from tonic firing in one cell of a coupled pair. A) TRN neurons fire bursts from hyperpolarized potentials (left; V_m_ = −82 mV) and tonic spikes from depolarized potentials (right; V_m_ = −55 mV). Scale bar 20 mV, 200 ms. B) Schematic of coupling measurement from an active cell (grey) and its electrically coupled neighbor (sage). C) Regular tonic spiking was driven by current injection into one cell (grey; V_m_ = −54 mV), while the coupled cell (sage) rests at −70 mV without stimulation. Experiments were performed with TTA in the bath. Scale bar 25 mV, 250 ms. D) Coupling coefficients (cc) for a pair before (left) and after (right) tonic spiking as shown in C. **cc** from the quiet cell into the active cell (green arrows) is shown in the top panel, and **cc** in the reverse direction on bottom. E) Coupling conductance G_C_ shown for each direction, before and after stimulated spiking, normalized to pre-activity values (for G_C_ quiet→ active, ΔG_C_ = 15.5 ± 2.4%, p_t_ = 0.004; for G_C_ active→ quiet, ΔG_C_ = −1.5 ± 0.5%, p_t_ = 0.78; n = 12 pairs). F) **cc** for each direction (for quiet→ active, Δcc = 13.6 ± 2.2%, p_t_ = 0.01, n = 12; for active → quiet, Δcc = 7.6 ± 0.5%, p_t_ = 0.018; n = 12 pairs). G) Input resistance for both cells during the experiments. H) G_C_ for each pair used in D (for G_C_ quiet→ active, p_s_ = 0.0009; for active → quiet, p_s_ = 0.76). I) **cc** for each pair used in E (for cc quiet→ active, p_s_ = 0.01; for active → quiet, p_s_ = 0.019). p_t_ indicates 2-tailed, paired Student’s t-test, and p_s_ is a Wilcoxon sign test.

LTP was specific to single-cell stimulation: paired activity in TTA did not significantly change synapse strength (Fig. 2A; p_t_ > 0.05 for both changes in cc and G_C_, averaged over both directions for paired activity, n = 8 pairs), and paired tonic spiking in unmodified ACSF also failed to change synapse strength (Fig. 2B; p_t_ > 0.05 for both changes in cc and G_C_, n = 6 pairs).

**Figure 2.**
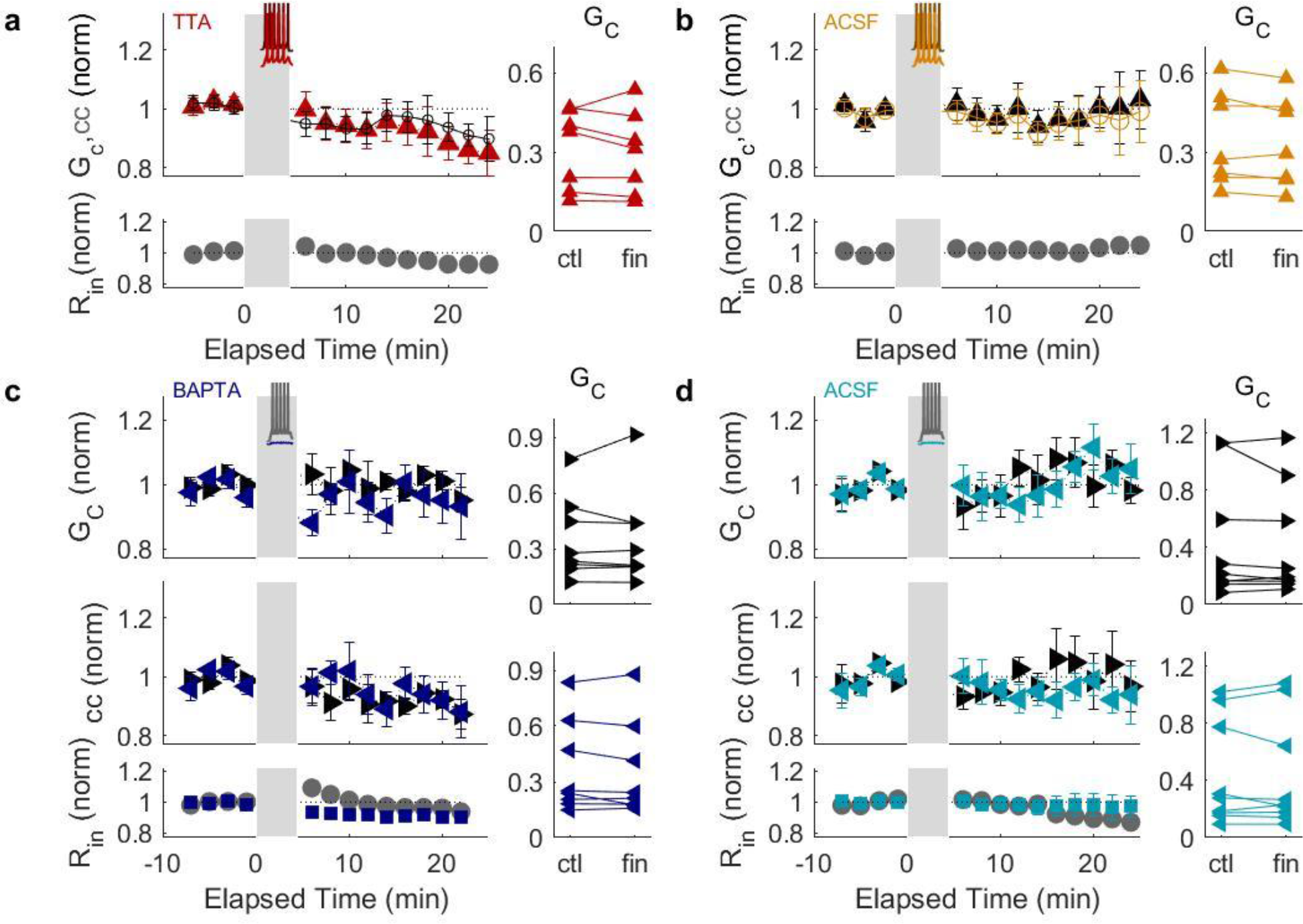
LTP is specific to single-cell stimulation and depends on calcium influx. A) cc and Gc before and after paired tonic spiking in TTA (averaged over both directions; for Δcc, p_t_ = 0.13, p_s_ = 0.12; for ΔG_C_, p_t_ = 0.3, p_s_ = 0.2; n = 8 pairs). B) cc and Gc before and after paired tonic spiking in unmodified ACSF (averaged over both directions; for Δcc, p_t_ = 0.37, p_s_ = 0.46; for ΔG_C_, p_t_ = 0.08, p_s_ = 0.07; n = 6 pairs). C) cc and Gc before and after single-cell spiking with calcium chelated by BAPTA in the internal. For ΔG_C_ active → quiet, p_t_ = 0.89, p_s_ = 0.84; for ΔG_C_ quiet → active, p_t_ = 0.34, p_s_ = 0.46; for Δcc active → quiet, p_t_ = 0.09, p_s_ = 0.07; for Δcc quiet → active, p_t_ = 0.66, p_s_ = 0.95 (n = 7 pairs). D) cc and Gc before and after single-cell tonic spiking in unmodified ACSF. For ΔG_C_ active → quiet, p_t_ = 0.38, p_s_ = 0.57; for ΔG_C_ quiet → active, p_t_ = 0.78, p_s_ = 0.82; for Δcc active → quiet, p_t_ = 0.4, p_s_ = 0.73; for Δcc quiet → active, p_t_ = 0.16, p_s_ = 0.25 (n = 9 pairs).

We hypothesized that the LTP we observed might be dependent on the smaller amounts of calcium influx from high-voltage activated channels (Budde et al., 1998). When we repeated single-cell tonic spiking in TTA with BAPTA in the pipettes to rapidly chelate all influxed calcium within both neurons, we observed no LTP (Fig. 2C; p_t_ > 0.05 for both directions of changes in cc or GC, n = 7 pairs). Single-cell spiking in unmodified ACSF also failed to induce changes in synaptic strength (Fig. 2D; p_t_ > 0.05 for both directions of changes in cc or G_C_, n = 9 pairs). Together, these results outline a calcium dependence of plasticity: minimal influx of calcium during single-cell tonic spiking, possibly buffered by the quiet cell, leads to LTP, while larger influx of calcium during paired tonic spiking activity exceeds that required for LTP, instead activating the mechanisms leading to LTD (Sevetson et al., 2017).

Our hypothesis that the LTP we saw arose from the minimal amount of calcium influx revealed by TTA and depolarization of the active cell led us to further investigate differences in calcium influx and transmission across the gap junction during the two different modes of induced activity. We therefore performed imaging experiments of pairs with the calcium indicator OGB-1 included in the internal solution (Fig. 3A_1_). We drove one cell to spike either in bursts or in tonic mode, while we held the quiet cell in held in voltage-clamp mode at −70 mV to minimize calcium signals arising from voltage-activated channels. During the induced bursting in one cell that leads to LTD (Haas et al., 2011), both peak and integrated calcium levels were higher in the active cell and in the quiet coupled neighbor than during induced tonic spiking (Fig. 3A_2_-A_4_). As a negative control, we repeated the imaging in a non-coupled pair (Fig. 3B_1-4_). Quantification of peak and integrated calcium signals in the quiet, coupled cells (Fig. 3C, D) both indicate that less calcium flows across the gap junction during tonic spiking than during bursting (for AUC, −25.2 ± 10.6%, p_s_ = 0.06; for peak calcium, −56.6 ± 10.9%, p_s_ = 0.03, n = 6 pairs).

**Figure 3.**
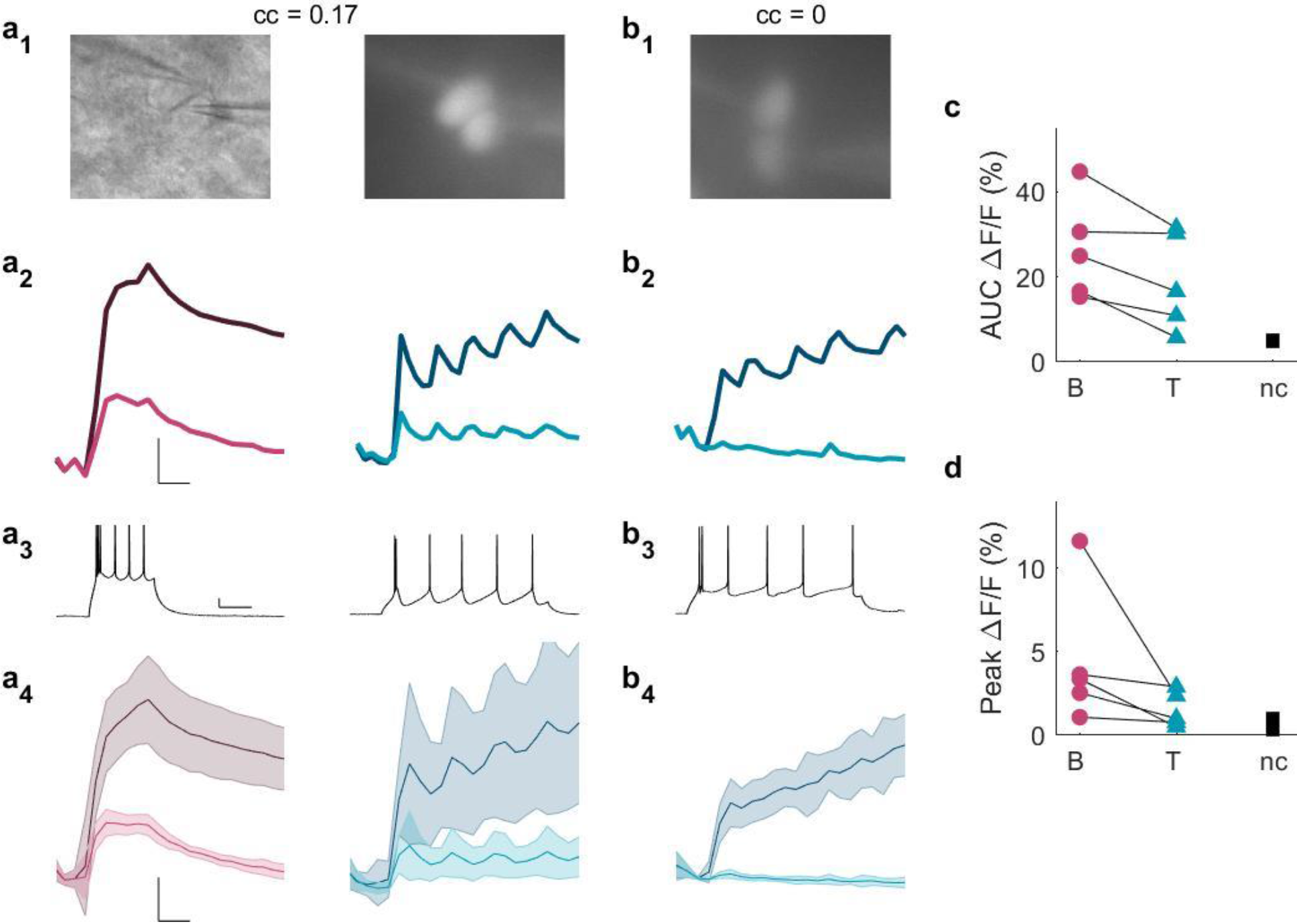
Calcium flows across the gap junction during spiking. A_1_) IR-DIC and GFP images of a coupled pair (cc = 0.17). A_2_) Calcium signals (ΔF/F) in a cell directly stimulated to burst (dark red) or spike tonically (dark blue), and calcium signals in the quiet cell (lighter shades) of a coupled pair. Scale bar 2.5%, 100 ms. A_3_) Bursting (left) or tonic spikes (center) that drove responses in A_2_. Scale bar 10 mV, 100 ms. A_4_) Average calcium signals during bursting (reds) and tonic spiking (blues) in this coupled pair. B_1_) GFP image of an uncoupled pair. Scale bar 2.5%, 100 ms. B_2_) Calcium signals (ΔF/F) in a cell directly stimulated spike tonically (dark blue), and in the quiet uncoupled cell (light blue). B_3_) Tonic spikes that drove responses in B_2_. B_4_) Average calcium signals during tonic spiking in uncoupled pairs (n = 2 pairs). C) Total (area under curve) ΔF/F in the quiet cell during bursting (B, red) and tonic spikes (T, blue) within each pair (mean difference −25.2 ± 10.6%, ps = 0.06, n = 6 pairs), and for uncoupled pairs (black, nc; n = 2 pairs). D) Peak ΔF/F in the quiet cell during bursting (B, red) and tonic spikes (T, blue) within each pair (mean difference −56.6 ± 10.9%, p_s_ = 0.03, n = 6 pairs), and for uncoupled pairs (black, nc; n = 2 pairs).

The LTP we observed that resulted from single-cell activity was expressed asymmetrically: coupling into the active cell increased, while coupling into the quiet cell decreased (Fig. 1). To examine whether this asymmetrical plasticity was consistent across pairs, we computed asymmetry as the ratio of coupling measured into the active cell, divided by coupling measured from the active cell. Those ratios consistently increased after LTP induction for both coupling coefficient (Fig. 4A; mean increase 24.3 ± 6.2%, p_t_ = 0.0015, p_s_ = 0.008, n = 12 pairs) and conductance (Fig. 4B: mean increase 22.4 ± 8.2%, p_t_ = 0.029, p_s_ = 0.008). We noted that some pairs went from initially symmetrical, to finally asymmetric, and *vice versa*. Changes in coupling asymmetry were uncorrelated with changes in R_in_ ratios (R^2^ = 0.39 for cc and 0.03 for G_C_). These results are consistent with the asymmetrical changes we also observed for burst-induced LTD (Haas et al., 2011) in that both asymmetrical LTP and LTD required sodium spikes, and changes were larger for coupling measured into the active cell. These changes in asymmetry indicate that the unidirectional use of the electrical synapse, and perhaps unidirectional calcium flow, systematically alter the fundamental property of each synapse.

**Figure 4.**
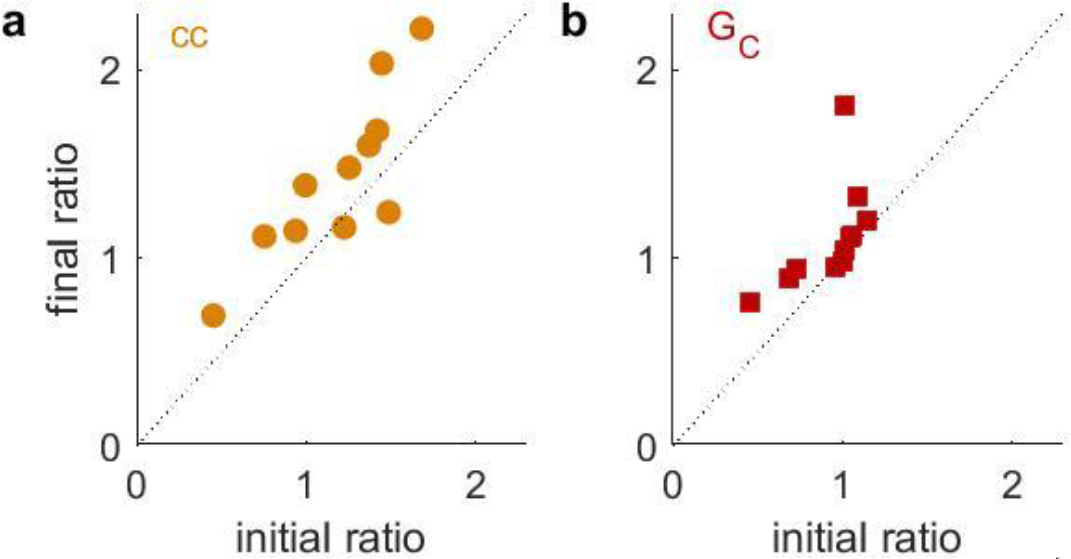
LTP is expressed asymmetrically. A) Asymmetry of coupling (**cc**into active cell / **cc** from active cell) increased consistently across pairs following activity-induced LTP, plotted here against initial values (mean change 24.3 ± 6.2%, p_t_ = 0.0015, p_s_ = 0.008, n = 12 pairs). B) Asymmetry of coupling conductance (G_C_ into active cell / G_C_ from active cell) increased consistently across pairs following activity-induced LTP, plotted against initial values (mean change 22.4 ± 8.2%, p_t_ = 0.029, p_s_ = 0.008).

Together, our results lead us to propose a calcium rule for plasticity of electrical synapses, whereby smaller amounts of calcium influx lead to LTP, and larger amounts lead to LTD (Fig. 5). The proposed rule is similar in concept to those proposed (Bienenstock et al., 1982; J. Lisman, 1989) (J. E. Lisman, 2001)and demonstrated (Dudek & Bear, 1992; Malenka & Bear, 2004; Malenka et al., 1989) at glutamatergic synapses.

**Figure 5.**
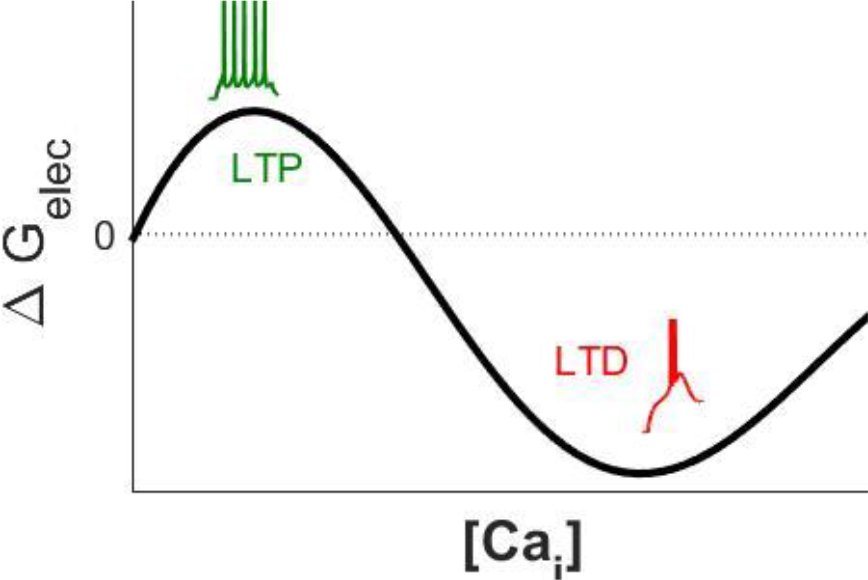
Proposed calcium-based activity rule for electrical synapses. Smaller influx of calcium during tonic spiking (green) leads to LTP, while larger influx during bursting (red) leads to LTD.

## Discussion

Here we show that long-term potentiation of electrical synapses results from tonic spiking in a single cell of the electrically coupled pair. To our knowledge, this is the first demonstration of LTP in mammalian electrical synapses that results from spiking activity within coupled neurons and flow of ions across the synapse. Moreover, our results taken together indicate a bidirectional calcium dependence of activity-dependent plasticity at electrical synapses, whereby high calcium influx and flow across the gap junction leads to LTD (Haas et al., 2011; Sevetson et al., 2017) while minimal calcium influx and flow across the gap junction leads to LTP. The links established here between physiological patterns of spiking activity in TRN neurons demonstrate that ongoing plasticity of electrical synapses is likely to occur in the active brain. As calcium rules are common for chemical synapses, we expect that calcium dependent rules, possibly in different forms, will underlie plasticity of electrical synapses across the brain.

The magnitude of LTP-induced changes in electrical synapse strength are similar in magnitude to our previous demonstration of LTD, less than 20% (GC or cc). These are modest changes relative to those often demonstrated at chemical synapses driven by tetanic stimulation. A direct comparison between synapse types and plasticity would need to account for differences in subcellular localization differences and differences in function, for instance spike efficacy. However, for LTD we showed that these changes were sufficient to silence a synapse, from one that induces spikes in neighbors, to an ineffective synapse (Haas et al., 2011). Further, these numerically modest changes in synaptic strength yield 5-10 ms changes in spike times in coupled neighbors (Haas, 2015). Computational models reinforce the effectiveness of changes in coupling in this range (Pham & Haas, 2018, 2019). Thus, even seemingly modest changes in electrical synapse strength produce physiologically substantial effects, and are poised to exert major influence on TRN synchrony and processing.

Our experiments required minimization of T currents by TTA in order to reveal conditions favorable for LTP at electrical synapses. While this is not strictly a physiological condition, we expect that together with depolarization of the active cell, these perturbations were necessary to counterbalance the artificial quiescence and hyperpolarization of the brain slice preparation, in which almost any depolarization of thalamic neurons activates T currents and thereby LTD. We expect that LTP is more likely to occur in vivo during prolonged depolarizations when tonic spiking dominates spiking patterns (Pinault, 2004). We further hypothesize that during tonic spiking in one neuron, calcium influx into the active cell is buffered across the gap junction by the quiet cell (Fig. 2, 3). Together, these results imply that LTP can be initiated by tonic spikes in single neurons, and subsequently counterbalanced by LTD resulting from bursts; together, these form a bidirectional basis for active neurons to modify the gain of their inputs.

Although functional asymmetry is not an expected property of gap junctions purely composed of connexin36 hemichannels (Srinivas et al., 1999), it has been observed as a widespread property of mammalian electrical synapses (Apostolides & Trussell, 2014; Devor & Yarom, 2002; Otsuka & Kawaguchi, 2013; Sevetson & Haas, 2015; Snipas et al., 2017; Vervaeke et al., 2010; Zolnik & Connors, 2016). Our previous results demonstrated that asymmetry systematically increases during LTD induced by asymmetrical bursting activity (Haas et al., 2011). Asymmetry was also shown to increase following LTP produced by NMDA application (Turecek et al., 2014) or by cerebellar input to coupled inferior olivary neurons (Lefler et al., 2014). Our results here add further evidence towards modifications of asymmetry that occur during plasticity; both our previous LTD results and the present LTP results indicate that sodium spikes are necessary for changes in asymmetry, and that changes are larger in the direction incoming to the active neuron. Together, these reports form a growing consensus that asymmetry and asymmetrical changes are a fundamental property of electrical synapses that potentially refine the function of each individual synapse within its neuronal circuit, allowing for each neuron to adjust the relative proportion of input it sends and/or receive from coupled neighbors via electrical synapses. Differences in intracellular scaffolding proteins have been shown to construct asymmetrical electrical synapses in zebrafish (Marsh et al., 2017). We suggest that increased expression or open probability of more-asymmetrical non-connexin36 proteins (Zolnik & Connors, 2016) could also account for this activity-dependent increase in asymmetry. Alternatively, differential post-translational modification, such as asymmetrical phosphorylation of cx36 hemichannels, or differences in ubiquitination-mediated endocytosis (Lynn et al., 2018), or the hypothesized effects of gating properties of the channel (Snipas et al., 2017) remain possible. In TRN, calcium bursts are activated in dendrites independently of somatic compartments, which could further result in independent modifications of one side of the gap junction. Asymmetry can strongly influence spike times in coupled pairs (Sevetson & Haas, 2015) and thereby impact cortical discrimination through the thalamocortical circuit (Pham & Haas, 2018). We suggest that it may also take part in directional signaling mediated by electrical synapses, such as that found in direction-sensitives retinal ganglion cells (Yao et al., 2018).

Based on our previous and current work, we propose a calcium rule for bidirectional activity-dependent plasticity at electrical synapses. This rule is ‘inverse’ to those described for chemical synapses, where smaller calcium influx produces LTD while larger influxes lead to LTP (Bienenstock et al., 1982; Dudek & Bear, 1992; Malenka et al., 1989). Notably, an inverse rule for chemical synapses also exists in the cerebellum (Coesmans et al., 2004). As previous studies have found that LTD is induced by calcium-based pathways leading to phosphatase activation (Sevetson et al., 2017), we expect that LTP may depend on a phosphorylation mechanism initiated by calcium. Retinal coupling depends on phosphorylation (Kothmann et al., 2007), and tetanus-induced forms of plasticity at mixed synapses onto Mauthner cells in goldfish depend on NMDA-regulated calcium entry (Pereda & Faber, 1996; Yang et al., 1990). Recent work has also shown that spiking-initiated calcium entry leads to potentiation for up to 10 min. at an invertebrate electrical synapse (Welzel & Schuster, 2018). While not directly examined, our results imply that the threshold between the concentration of calcium for LTP and LTD is rather low, as even paired tonic spiking fails to induce LTP, and the T current that underlies burst-induced LTD is active at rest potentials. This bias of plasticity towards LTD, however, could be specific to brain-slice conditions, as addressed above. We find it likely that electrical synapses across the brain follow a calcium-based rule for plasticity, although non-bursting neuronal types may alternatively follow a different rule.

A role for plasticity of electrical synapses has been proposed for switches in attentional state (Coulon & Landisman, 2017). Taken together, our results demonstrate that the strength of electrical synapses can be modified positively or negatively in strength as a result of physiological activity in the TRN, and with neuron-specific directionality. For single cells, activity-dependent plasticity of electrical synapses could modulate that cell’s input sensitivity between modes in which intra-TRN input is strongest and dominates its responses, to modes in which corticothalamic or thalamocortical chemical input is given preference over electrically networked signals. This high degree of acuity and adjustability of connectivity within coupled networks forms a basis for shifts of sensory attention regulated by the TRN. TRN neurons can toggle thalamic neurons between firing modes in order to maintain cortical sleep rhythms and the associated behaviors of that state (Sorokin et al., 2017), and the activity and synchrony of TRN neurons required for that toggle may be supplied by its electrical synapses. Sensory processing of selective features (Soto-Sanchez et al., 2017) is also likely to depend on acutely refined TRN connectivity in a similar manner. Beyond the TRN and its functions, our work shows that electrically coupled networks are potentially under a high degree of regulation of function throughout the mammalian brain.

## Methods

### Electrophysiology

All experiments were performed in accordance with federal and Lehigh University IACUC animal welfare guidelines. Sprague-Dawley rats of both sexes aged postnatal day 11-15 were anesthetized by inhaled isoflurane (5 mL of isoflurane applied to fabric, within a 1 L chamber) and sacrificed via decapitation. Horizontal brain slices 300-400 μm thick were cut and incubated in sucrose solution (in mM): 72 sucrose, 83 NaCl, 2.5KCl, 1 NaPO_4_, 3.3 MgSO_4_, 26.2 NaHCO_3_, 22 dextrose, 0.5 CaCl_2_. Slices were incubated at 37°C for 20 min following cutting and returned to room temperature until recording. The bath for solution during recording contained (in mM): 126 NaCl, 3 KCl, 1.25 NaH_2_PO_4_, 2 MgSO_4_, 26 NaHCO_3_, 10 dextrose and 2 CaCl_2_, 300–305 mOsm L−1, saturated with 95% O_2_/5% CO_2_. The submersion recording chamber was held at 34°C (TC-324B, Warner Instruments). Micropipettes were filled with (in mM): 135 potassium gluconate, 2 KCl, 4 NaCl, 10 Hepes, 0.2 EGTA, 4 ATP-Mg, 0.3 GTP-Tris, and 10 phosphocreatine-Tris (pH 7.25, 295 mOsm L^−1^). 1 M KOH was used to adjust pH of the internal solution. Internal containing 1,2-bis(2-aminophenoxy)ethane-N,N,N’, N’-tetraacetic acid (BAPTA) had concentrations of 10 μM. The approximate bath flowrate was 2 ml min^−1^ and the recording chamber held approximately 5 ml solution. The specific T-channel antagonist TTA-A2, generously provided by Dr Bruce Bean (Harvard University) or TTA-P2 (Alamone) were made into stock aliquots of 3 mM in DMSO; final concentration was 1 μM. 6-Cyano-7-nitroquinoxaline-2,3-dione (CNQX) at 2.5 μM was obtained from Sigma (St. Louis, MO, USA), and diluted into high-concentration stock solutions in DMSO or water before final dilution. Final DMSO concentration was always <0.2%.

The TRN was visualized under 5x magnification, and pairs of TRN cells were identified and patched under 40× IR-DIC optics (SliceScope, Scientifica, Uckfield, UK). Voltage signals were amplified and low-pass filtered at 8 kHz (MultiClamp, Axon Instruments, Molecular Devices, Sunnyvale, CA, USA), digitized at 20 kHz (custom Matlab routines controlling a National Instruments (Austin, TX, USA) USB6221 DAQ board), and data were stored for offline analysis in Matlab (Mathworks, R2017a, Natick, MA, USA). Recordings were made in whole-cell current-clamp mode. Values V_rest_ ranged from −50 to −70 mV. 500-ms pulses of negative injections of current were used to measure coupling, with amplitudes of injected current minimized in order to minimize T current activation in the injected cell, with a goal of 0.5 - 1 mV deflection in the coupled cell. Pipette resistances were 5-9 MΩ before bridge balance, which was removed if exceeding 25 MΩ. Voltages are reported uncorrected for the liquid junction potential.

### Calcium imaging

To visualize calcium flux into and within coupled and uncoupled cells, 200 μM Oregon Green 488 BAPTA-1 (OGB-1) was added to the internal solution. An LED with wavelength 472 nm was delivered through the objective to excite the OGB-1. Images were captured at 30 fps through 40X IR-DIC optics (SliceScope, Scientifica, Uckfield, UK) and stored for offline analysis with Matlab. Regions of interest (ROIs) were chosen in the center of each recorded neuron, and changes in fluorescence normalized to baseline and background fluorescence were computed for each ROI.

### Numerical Analysis

All numerical analysis was performed in Matlab (R2017). Input resistances (R_in_) for each cell and coupling between cells were quantified by injecting 25-100 pA of hyperpolarizing current into one cell of a coupled pair and measuring the voltage deflection in that cell (ΔV) and in the couple neighbor (δV). Coupling coefficient (cc) is computed as δV/ΔV and are reported as averages of a set of 10 measurements, repeated every 2 min. Coupling conductances G_C_ were estimated separately for each direction (Fortier, 2010; Sevetson & Haas, 2015). Experiments were discarded if input resistance R_in_ of either cell deviated from its initial value by more than 20%. Changes in coupling were evaluated as the average over the first 20 minutes following activity, compared to the normalized baseline values, and are reported as means ± SEM. We report Student’s *t*-tests as two-tailed paired comparisons of pre- and post-stimulus averages and report the results as p_t_. Two-sided Wilcoxon signed rank tests were also carried out on the sets of change in coupling for each condition and are reported as p_s_. No multiple comparisons were performed.

## Acknowledgements

Bruce Bean (Harvard) generously supplied TTA-A2. We thank C. E. Landisman, R. M. Burger, T. Pham and all Haas lab members for input on drafts of this manuscript. Funding was provided by the NSF (IOS 1557474), the Whitehall Foundation, and the Brain & Behavior Foundation.

